# Fmn2 regulates growth cone motility by mediating a molecular clutch to generate traction forces

**DOI:** 10.1101/2020.02.21.959759

**Authors:** Ketakee Ghate, Sampada P. Mutalik, Lakshmi Kavitha Sthanam, Shamik Sen, Aurnab Ghose

## Abstract

Growth cone - mediated axonal outgrowth and accurate synaptic targeting are central to brain morphogenesis. Translocation of the growth cone necessitates mechanochemical regulation of cell - extracellular matrix interactions and the generation of propulsive traction forces onto the growth environment. However, the molecular mechanisms subserving force generation by growth cones remain poorly characterized. The formin family member, Fmn2, has been identified earlier as a regulator of growth cone motility. Here, we explore the mechanisms underlying Fmn2 function in the growth cone. Evaluation of multiple components of the adhesion complexes suggests that Fmn2 regulates point contact stability. Analysis of F-actin retrograde flow reveals that Fmn2 functions as a clutch molecule and mediates the coupling of the actin cytoskeleton to the growth substrate, via point contact adhesion complexes. Using traction force microscopy, we show that the Fmn2-mediated clutch function is necessary for the generation of traction stresses by neurons. Our findings suggest that Fmn2, a protein associated with neurodevelopmental and neurodegenerative disorders, is a key regulator of a molecular clutch activity and consequently motility of neuronal growth cones.

## INTRODUCTION

Neurite outgrowth and pathfinding is critical for the establishment of neural circuits during development and also for the re-establishment of neural tracts following injury. Axon guidance involves directed motility of the highly dynamic chemosensory growth cones. Like the migration of non-neuronal cells, motility of growth cones involves precise coordination between the sensing of environmental mechanochemical cues and directed remodelling of the underlying cytoskeleton.

Neuronal growth cones instructively remodel their cytoskeleton to generate protrusive processes (like filopodia and lamellipodial veils), establish directional polarity and develop contractility that is transmitted as traction forces onto the growth substrate (Lowery and Vactor 2009; Kerstein *et al*. 2015). Cell surface receptors, like the integrins, engage with extracellular matrix (ECM) molecules while templating adhesion complexes intracellularly that facilitate physical coupling with the force-generating actomyosin cytoskeleton (Mitchison and Kirschner 1988; Swaminathan and Waterman 2016). Actin polymerization and myosin activity result in centripetally oriented F-actin retrograde flow (Mitchison and Kirschner 1988; Medeiros *et al*. 2006). Adhesion complexes, known as point contacts, form a regulatable physical coupling between the F-actin retrograde flow and the growth substrate. This coupling facilitates the transmission of the forces generated by actomyosin activity onto the substrate in the form of traction stresses allowing forward propulsion of the cell. The variable coupling of the actin cytoskeleton to adhesion sites has been described as a molecular clutch (Mitchison and Kirschner 1988; Swaminathan and Waterman 2016). In growth cones, adhesion to ECM molecules is known to engage the clutch resulting in a reduction in F-actin retrograde flow rates at the sites of adhesion (Santiago-Medina *et al*. 2013; Swaminathan and Waterman 2016). In this model, the strength of the clutch activity (the ability to couple with the actomyosin system) directly regulates the translocation rates. Further, local reduction in F-actin retrograde flow also remodels the microtubule cytoskeleton and facilitates directional outgrowth (Schaefer *et al*. 2008). The molecular clutch can be modulated by substrate stiffness, allowing it to generate traction in stiffness dependent manner. On stiffer substrates, the clutch engages over a short time resulting in frictional slippage whereas on compliant substrates, the engagement is more long-lasting with sudden intermittent detachments (Chan and Odde 2008).

Unlike in non-neuronal cells, relatively fewer studies have directly explored the molecular clutch model in growth cones and identified clutch molecules. F-actin retrograde flow was reduced at the adhesion sites in *Aplysia* growth cones responding to adhesive substrates (Lin and Forscher 1995; Suter and Forscher 2000; Suter and Forscher 2001; Lee and Suter 2008). Several adaptor and signalling proteins are recruited to the adhesive point contacts and possibly facilitate the clutch function and its regulation (Bard *et al*. 2008; Myers and Gomez 2011; Toriyama *et al*. 2013). The molecular clutch model in growth cones is underscored by studies demonstrating local modulation of retrograde flow at point contacts (Santiago-Medina *et al*. 2013) and the association of point contact density with the rate of F-actin retrograde flow (Koch *et al*. 2012). Due to the clutching activity, the point contacts develop traction stresses on the growth substrate and local variations in the magnitude of these forces dominate the directionality of the extending growth cone (Suter and Forscher 2001; Chan and Odde 2008; Buck *et al*. 2016). Cue-dependent localized remodelling of the point contacts (Moore *et al*. 2009; Moore *et al*. 2012; Baba *et al*. 2018) resulting in differential clutching activity is thought to contribute to growth cone turning during haptotaxis. While growth cone filopodia are themselves contractile and generate traction stresses, the forces developed by the point contacts in the lamellipodia and transitional zone of the growth overwhelmingly contribute to the overall growth cone traction (Betz *et al*. 2011; Hyland *et al*. 2014). Consequently, loss of filopodia, while resulting in loss of directionality, does not have a strong effect of translocation (Marsh and Letourneau 1984). Thus identifying the regulation of the force-generating clutch mechanism in the growth cone is central to our understanding of growth cone-mediated axonal outgrowth.

Fmn2, a non-Diaphanous related formin, has been implicated in cognitive dysfunction (Almuqbil *et al*. 2013; Law *et al*. 2014) and de-regulation of sensory processing in humans (Marco *et al*. 2018). In post-traumatic stress disorder and Alzheimer’s disease patients, Fmn2 expression is reduced (Agís-Balboa *et al*. 2017). Fmn2 knockout mice show compromised associative memory and accelerated age-associated memory decline (Peleg *et al*. 2010; Agís-Balboa *et al*. 2017). However, the mechanistic basis of Fmn2 function in neurons is poorly characterized. We have previously shown that Fmn2 regulates the stability of growth cone filopodia in spinal commissural neurons and pathfinding of these neurons *in vivo* (Sahasrabudhe *et al*. 2016). However, this study did not explain the mechanism underlying reduced growth cone translocation rates. Here we identify the contribution of Fmn2 in stabilising growth cone adhesions and generating traction stresses underlying motility. We find that Fmn2 acts as a clutch molecule and regulates the stability of the adhesive point contacts and their ability to generate traction stresses on the substrate.

## MATERIALS AND METHODS

### Transfection, culture, immunostaining and imaging of primary neurons

Six-day old chicken eggs (*Gallus gallus*; HH stage 25/26) were utilised to harvest entire spinal cords of embryos. The embryos were dissected in sterile phosphate buffered saline (PBS) on a Sylgard-coated plate under a dissecting microscope inside a horizontal laminar flow hood. The dissected tissue was collected in embryonic medium (Leibovitz’s media (Gibco) with 1X penicillin-streptomycin (Gibco)). The spinal cord was spun at 3000 rpm for 3 min at room temperature after which the embryonic medium was replaced with 1x trypsin-EDTA (Lonza). The tissue was macerated in trypsin and incubated at 37°C for 15-20 mins. The further embryonic medium was added to the trypsin and the dissociated spinal cord was spun at 3000 rpm for 3 min. The cells were plated in 2 ml of basal culture medium (L-15 (Gibco), 1X PenStrep (Gibco), 10% heat-inactivated FBS (Gibco)) with 20ng/ml NGF (Invitrogen)) and incubated without CO_2_ at 37°C for 24-36 hrs.

The control morpholino (Control-MO) sequence used as a negative control: CCTCTTACCTCAGTTACAATTTATA, the anti-chick translation-blocking Fmn2 morpholino (Fmn2-MO) sequence used: CCATCTTGATTCCCCATGATTTTTC. The Fmn2-MO has been previously extensively validated for efficacy and specificity with the evaluation of Fmn2 protein levels and rescue experiments both *in vivo* and *in vitro* (Sahasrabudhe *et al*. 2016). This study also reports a significant reduction in endogenous Fmn2 protein and functional rescue with morpholino-resistant Fmn2 cDNA.

For electroporation, cells were resuspended in 100 μl OptiMEM (Gibco) with 10 μg of plasmid and 100 μM morpholino. The cell suspension with plasmid and/or morpholinos was transferred to the electroporation cuvette and current was delivered in form of five 20 V pulses where each pulse lasted for 50ms with an inter-pulse interval of 100ms using NEPA-21 electroporator (Nepagene). Following electroporation, the cell suspension was transferred to a microfuge tube with 400 μl OptiMEM (Gibco) and finally plated on 20 ng/ml fibronectin (Sigma-Aldrich) coated coverslip bottom plates with 2 ml of basal culture media (mentioned above).

### Live imaging and analysis of neuronal growth cone motility

Around thirty three hours post electroporation the samples were imaged using a PlanApoN 60×/1.49 oil immersion objective on an Olympus IX81 system equipped with a Hamamatsu ORCA-R2 CCD camera. Images were captured every 10 s for up to 30 min. The system was equipped with a focus drift correction mechanism and the imaging was conducted at 37°C without CO_2_. The growth cone was identified as a dynamic, often fan shaped structure at the distal end of the axon to assess the motility parameters of which such as accumulated distance, speed and directionality, a substack of 60 timepoints was made from the original movie of 30 min (180 frames). Each frame of the substack has a time interval of 30 seconds. The growth cone boundary was manually drawn using the GFP signal on each frame and the region of interests (ROIs) were saved in the ROI manager in Fiji. A sharp narrowing of the growth cone was used to demarcate it from the axon. In case of any ambiguity in the specific frame, immediately preceding and following frames were evaluated to aid the identification of the growth cone ‘neck’. The investigators were blinded to the genotypes while generating the ROIs.

To measure the centroid of the growth cone, a new blank hyperstack of 61 frames similar to the original time-lapse was created on which each ROI of the growth cone was overlaid and the centroid identified. The x and y coordinates of the centroid were used to measure accumulated distance, speed and directionality of the growth cone.

In order to calculate the distance between two coordinates following formula was used: 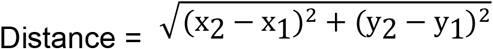, where (x_1_, y_1_) and (x_2_, y_2_) are coordinates of two successive timepoints. For measuring accumulated distance, the distance between every consecutive centroid was calculated and added together. Velocity was obtained by dividing this distance with the total time of imaging (30 min) and directionality was acquired by determining the ratio of displacement (distance between centroid of the first and last frame) to the total accumulated distance.

### Immunostaining, imaging and analysis of growth cone point contacts

Spinal neurons were electroporated and cultured with morpholino and pCAG-GFP (Addgene) as described earlier. After 24 – 30 hrs the cells were fixed using modified Kreb’s fixative (Dent and Meiri 1992). 50% Kreb’s buffer (145mM NaCl, 5mM KCl, 1.2mM CaCl_2_, 1.3mM MgCl_2_, 1.2mM NaH_2_PO_4_, 10mM glucose, 20mM HEPES) with 0.4M sucrose was mixed with 50% Bouin’s fixative (75 ml Saturated Picric acid, 20 ml Formaldehyde (37-40 %) and 5 ml glacial acetic acid) and cells were fixed in this buffer for 15 mins at room temperature. Cells were further permeabilized for 20 mins in 0.01% saponin in 10% heat-inactivated normal goat serum in PBS. Following permeabilization, the samples were washed in 0.5% BSA in Kreb’s buffer twice for 10 min each. Further, cells were blocked in 3% BSA in PBS for 1 hr at room temperature. Antibodies against total FAK (Abcam, ab33141), phosphor-FAK-Y397 (Abcam, ab4803), Vinculin (Abcam, ab11194-200), total Pax (Abcam, ab2264), phosphor-Pax-Y31 (Abcam, ab4832) were added in blocking buffer at 1:600 dilution and the cells were incubated overnight at 4°C.

After removing primary antibodies, the cells were washed twice in 0.5% BSA in Kreb’s buffer for 10 min each. The samples were blocked with 10% normal goat serum in PBS for 45-60 min and then 1:300 dilution of secondary antibody (Invitrogen) was added and incubated for 60 min at room temperature. After the secondary antibody, the cells were washed thrice in 0.5% BSA in Kreb’s buffer for 10 min each. Samples were finally mounted in 80% glycerol.

Growth cone point contacts were imaged using a Zeiss 710 inverted confocal microscope with PlanApoN 63x/1.40 oil immersion objective. The images were acquired with 2x zoom and all imaging conditions were kept constant across treatments and samples so that fluorescence intensities could be compared.

For analysing point contacts, the images were manually processed in Fiji. The growth cone was demarcated using the GFP signal by investigators blind to the genotypes. Only growth cones with clearly identifiable ‘necks’ were chosen for analysis. Analysis of point contacts was limited to the area defined by the outline of the growth cone. The antibody signal was subjected to background subtraction to reduce the noise followed by application of unsharp mask which sharpened the boundaries of the point contact staining observed. These images were thresholded using the Otsu algorithm and the resultant binary image was processed through the particle analyser with 1.5 pixels as a baseline threshold to obtain a mask demarcating the outlines around point contacts within the selected growth cone area. This mask could be overlayed on the raw image of point contact staining to get extract the original signal intensity of the point contacts.

The particle analysis tool in Fiji generated the number of point contacts detected and the addition of the mask on the antibody channel yielded the intensity at each detected point contacts. These outputs from Fiji were exported to Microsoft Excel 2007 and plotted using GraphPad Prism 5 for further analysis.

### Evaluating F-actin retrograde flow

Neurons transfected with morpholinos and pCAG mGFP-Actin (Addgene #21948) were cultured on the fibronectin-coated glass and imaged after 24 hrs using a PlanApoN 100x/1.49 oil immersion objective on an Olympus IX81 system equipped with a Hamamatsu ORCA-R2 CCD camera. Images were captured every 1 s for up to 3 min. The system was equipped with a focus drift correction mechanism and the imaging was conducted at 37°C without CO_2_.

Following generation of substacks of the original movie in Fiji, kymographs were generated using the MetaMorph software (Molecular Devices) using a segmented line tool (width: 5 pixels). Velocities were calculated from the kymographs using the FlowTrack code (obtained from Dr. D. Odde, University of Minnesota; (Chan and Odde 2008) in the Matlab 2007b. Retrogradely moving regions from the heat map generated of the kymograph were selected using a rectangular selection. The analysis was restricted to sections of the time-lapse series where the filopodia were attached and not dynamic.

### Traction force microscopy

Filopodial traction force assays were modified from an earlier protocol (Chan and Odde 2008). 28 mm clean coverslips were treated with 0.5% APMS-3-aminopropyltrimethoxysilane (APMS) (Sigma) in water for 15-20 min followed by 4-5 washes with distilled water. Treated coverslips were baked at 160°C for 1-2 hrs. APMS treated coverslips were attached to 35 mm petri dishes with holes drilled at the bottom using silicone glue with the APMS treated side facing inside the dish. The glue was allowed to dry overnight, following which 0.5% glutaraldehyde (Sigma) was added onto the coverslips and incubated for 1 hr at room temperature. Further, the coverslips were washed with water 3-4 times and allowed to dry completely. Polyacrylamide (PAA) gel mixture for stiffness 0.438 ± 0.064 kPa was prepared using 3% acrylamide and 0.1% bisacrylamide as described previously in (Tse and Engler 2010) and 15 μl of the gel mixture was added onto each of the coverslip bottom culture plates. To transfer fluorescent beads onto the gels, 18 mm clean coverslips were incubated with 50 μg/ml fibronectin (Sigma) at 37°C for 30 mins. After removing the fibronectin solution, the coverslips were allowed to dry completely. 100 μl of 1:20 diluted Fluorosphere carboxylate modified beads (Thermo) (200 nm) from 1:1 working stock was spin-coated on these coverslips at 1500 rpm for 5 min.

Immediately after adding 15 μl of the gel mixture onto the treated coverslip attached to the petri dish, the smaller bead coated coverslip was inverted on the gel with the bead-coated side facing the gel mixture. Once the gel was solidified, MiliQ water was used to keep the gels hydrated and the top coverslips were carefully removed. To cross-link ECM substrate of the PAA gels, 200 μl of 2 mg/ml sulfo-SANPAH (Covachem) was added to the gels and allowed to coat the gels for 1 hr at 5 to 8 cm distance from a portable UV lamp (365 nm) inside a cell culture hood. The plates were later washed with sterile distilled water 3 times and incubated with 250 μl of 50 μg/ml fibronectin (Sigma) overnight at 4°C until further neuronal culturing.

Stiffness of the gels prepared was measured using atomic force microscopy (AFM). Two technical replicates, each with minimum 30 readings were performed using a pyramidal cantilever. The gel stiffness was estimated to be 0.438 ± 0.064 kPa. Data analysis was done using Hertz fitting (MacKay and Kumar 2012).

The growth cones were imaged with PlanApoN 100x/1.49 oil immersion objective on an Olympus IX81 system equipped with a Hamamatsu ORCA-R2 CCD camera and PlanApoN 60x/1.42 oil immersion objective on an Olympus IX83 system equipped with a regular CCD camera after 48 hrs of culture. An image of the growth cone and corresponding image of the bead distribution were captured. This was the stressed image. Trypsin (10x, Sigma-Aldrich) was added to the plate with 5x final concentration. After trypsin flow, and following complete detachment of the growth cone, an image of the unstressed bead distribution was captured again using similar settings. For the traction force analysis, the images of beads pre and post trypsin deadhesion were stabilized using a template matching plugin in Fiji. The images were analysed using the Fourier Transform Traction Cytometry (FTTC) method established by Butler and colleagues (Butler *et al*. 2002). The following parameters were used: gel stiffness (0.438 kPa), Poisson ratio for the gel (0.48) and gel thickness (60 μm). The code generates displacement fields by employing the cross-correlation between stressed and relaxed images after regularization and further generates the traction force values.

### Data representation and statistics

All data were plotted and statistically analysed in GraphPad Prism 5. Box and Whisker plots represent the spread of the data using the Tukey method. The bottom and top of the box are the first and third quartiles, the whiskers span the lowest datum within the 1.5 interquartile range of the lower quartile and the highest datum within the 1.5 interquartile range of the upper quartile. Outliers are represented outside the box as individual data points. In the case of bar graphs, error bars represent the standard error of the mean (SEM). Mann-Whitney test in GraphPad Prism 5 was used to compare the data.

## RESULTS

### Fmn2 regulates growth cone motility

Role of Fmn2 in axonal pathfinding *in vivo* and its regulation of filopodial stability has been demonstrated in chick spinal neurons (Sahasrabudhe *et al*. 2016). While this study reported deficits in the outgrowth of spinal neurons, the underlying mechanism was not investigated. To identify the molecular mechanisms underlying Fmn2-dependent regulation of growth cone motility, we evaluated growth cone dynamics of chick spinal neurons plated on fibronectin-coated coverglass. Neurons were transfected either with a morpholino optimised to knockdown Fmn2 (Fmn2-MO) or a control morpholino (Control-MO) along with soluble GFP to detect transfected neurons. Only transfected neurons were considered for live imaging and analysis (Figure 1A-C; Movie S1). Using antibodies against endogenous Fmn2, we found that at least 70% knockdown was achieved by morpholino treatment (Figure S1). Live imaging revealed that the accumulated distance of the Fmn2 knockdown growth cones was significantly reduced (Figure 1D; 52.40 ± 4.037 μm) compared to controls (Figure 1D; 68.84 ± 4.295 μm). Additionally, Fmn2-depleted growth cones, though dynamic, translocated slowly (Figure 1E; 1.747 ± 0.135 μm/min) compared to Control-MO transfected neurons (Figure 1E; 2.295 ± 0.143 μm/min). The persistent directionality of movement (the ratio of the displacement to the total accumulated distance covered) was also compromised upon Fmn2 knockdown (Figure 1F; Fmn2-MO: 0.0516 ± 0.01; Control-MO: 0.1589 ± 0.023). Co-expressing mouse Fmn2 cDNA (mFmn2), which is resistant to the morpholino, rescued all the motility parameters (Figure 1D-F).

**Figure 1:**
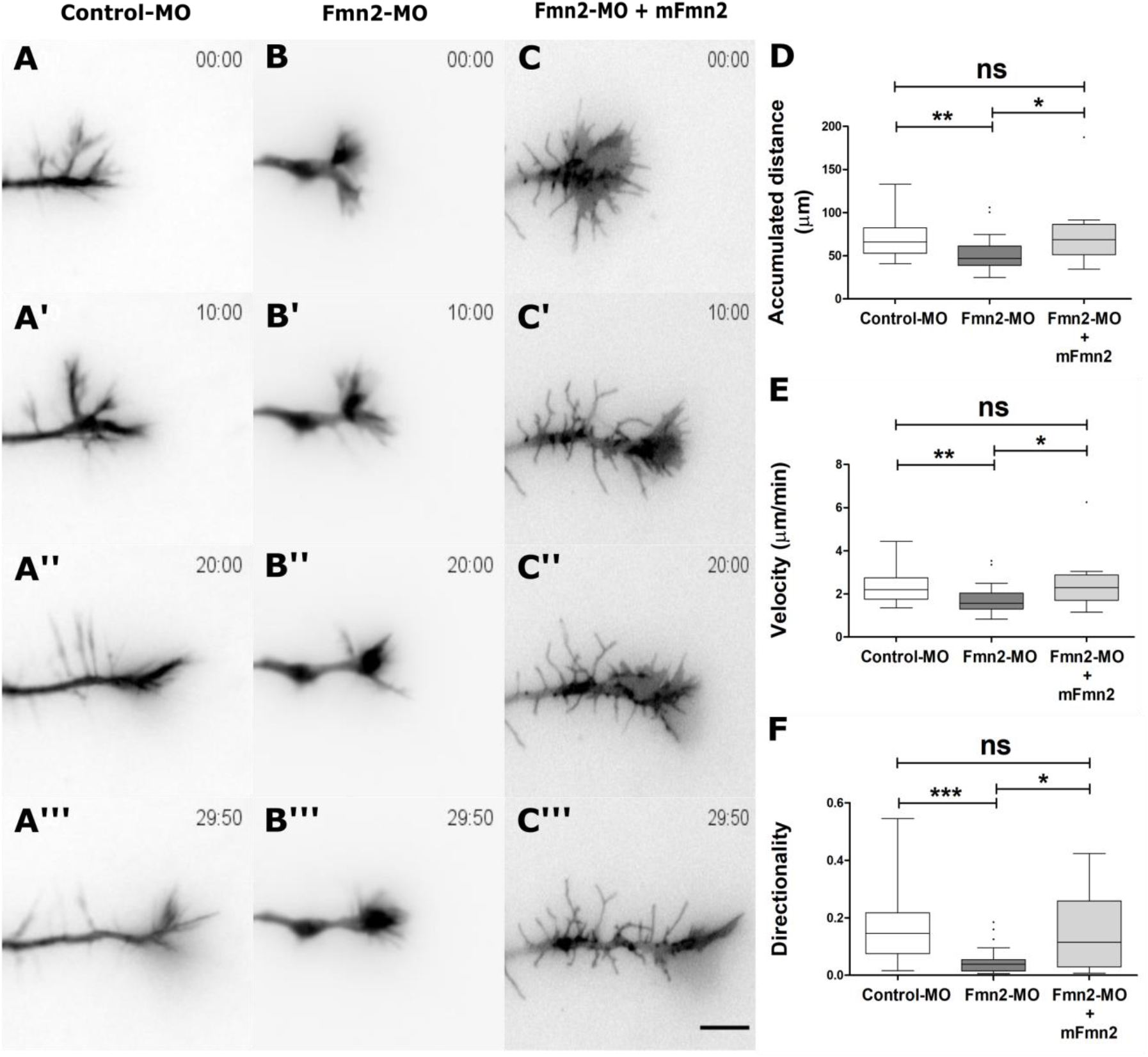
Fmn2 regulates growth cone motility. (A – A”‘) Representative snapshots from time-lapse movie of spinal growth cone transfected with Control morpholino (Control-MO) expressing cytoplasmic GFP. (B – B’’’) Representative snapshots from time-lapse movie of spinal growth cone transfected with Fmn2 morpholino (Fmn2-MO) expressing cytoplasmic GFP. (C – C’’’) Representative snapshots from time-lapse movie of spinal growth cone transfected with Fmn2 morpholino (Fmn2-MO) along with morpholino-resistant mouse Fmn2 cDNA (mFmn2). The growth cones were imaged for a total period of 30 minutes to quantify the motility parameters. (D) Graph comparing accumulated distance of Fmn2 morphant growth cones compared to the control growth cones, however, this phenotype is rescued in growth cones co-transfected with morpholino-resistant mFmn2. (E) Graph showing reduced translocation rates (accumulated distance divided by total time of imaging) of Fmn2 depleted growth cones compared to, this phenotype is rescued in growth cones expressing mFmn2. (F) Quantitation of directionality shows a reduction upon Fmn2 knockdown in spinal growth cones which can be rescued by mFmn2. The number of growth cones analysed for Control-MO, Fmn2-MO and rescue experiments are 23, 24 and 16, respectively. Data are represented as box and whisker plots using the Tukey method. The horizontal line inside the box represents the median. Statistical comparisons were performed using the Mann-Whitney test; *, p≤0.05, **, p≤0.01, ***, p≤0.001, ns, non-significant. Scale bar, 5 μm in all images.

These experiments underscored the function of Fmn2 in regulating growth cone-mediated motility of spinal neurons.

### Fmn2 mediates point contact stability in growth cones

Cellular motility is a complex process achieved by the coordinated activities of cytoskeletal remodelling, dynamic regulation of adhesion complexes with the substrate and force generation by actomyosin activity. We have previously reported that Fmn2 regulates the stability of point contacts in growth cone filopodia and focal adhesion stability in fibroblasts (Sahasrabudhe *et al*. 2016). Fmn2 depletion resulted in the reduction of tyrosine 397 phosphorylated focal adhesion kinase (pFAK; a marker of adhesion complex stability) signal in the entire growth cone (Sahasrabudhe *et al*. 2016). These observations prompted us to explore in detail the regulation of growth cone - extracellular matrix (ECM) adhesion complexes by Fmn2. Using antibodies against several constituents of growth cone point contacts, we undertook a detailed evaluation of the distribution and status of these adhesion complexes.

Spinal neurons transfected with morpholinos and a soluble GFP reporter plasmid were cultured on fibronectin-coated glass for 24 hrs after which the neurons were fixed using Kreb’s fixative (Dent and Meiri 1992) and immunostained with antibodies against Paxillin, phospho-Paxillin (Y31), Vinculin, Focal adhesion kinase (FAK) and phospho-FAK (Y397). The growth cones were imaged using confocal microscopy and multiple features of the point contacts were evaluated (Figure 2A-J). The growth cone area outlines were marked manually using the GFP signal. The images were thresholded using the Otsu algorithm and processed to generate a mask demarcating the area occupied by point contacts. This approach not only allowed us to score the number of individual point contacts but could also be overlaid on the original image to extract the fluorescence signal intensity within the demarcated area. This enabled us to compare the changes in the endogenous levels of the point contact proteins between control and Fmn2-depleted growth cones.

**Figure 2:**
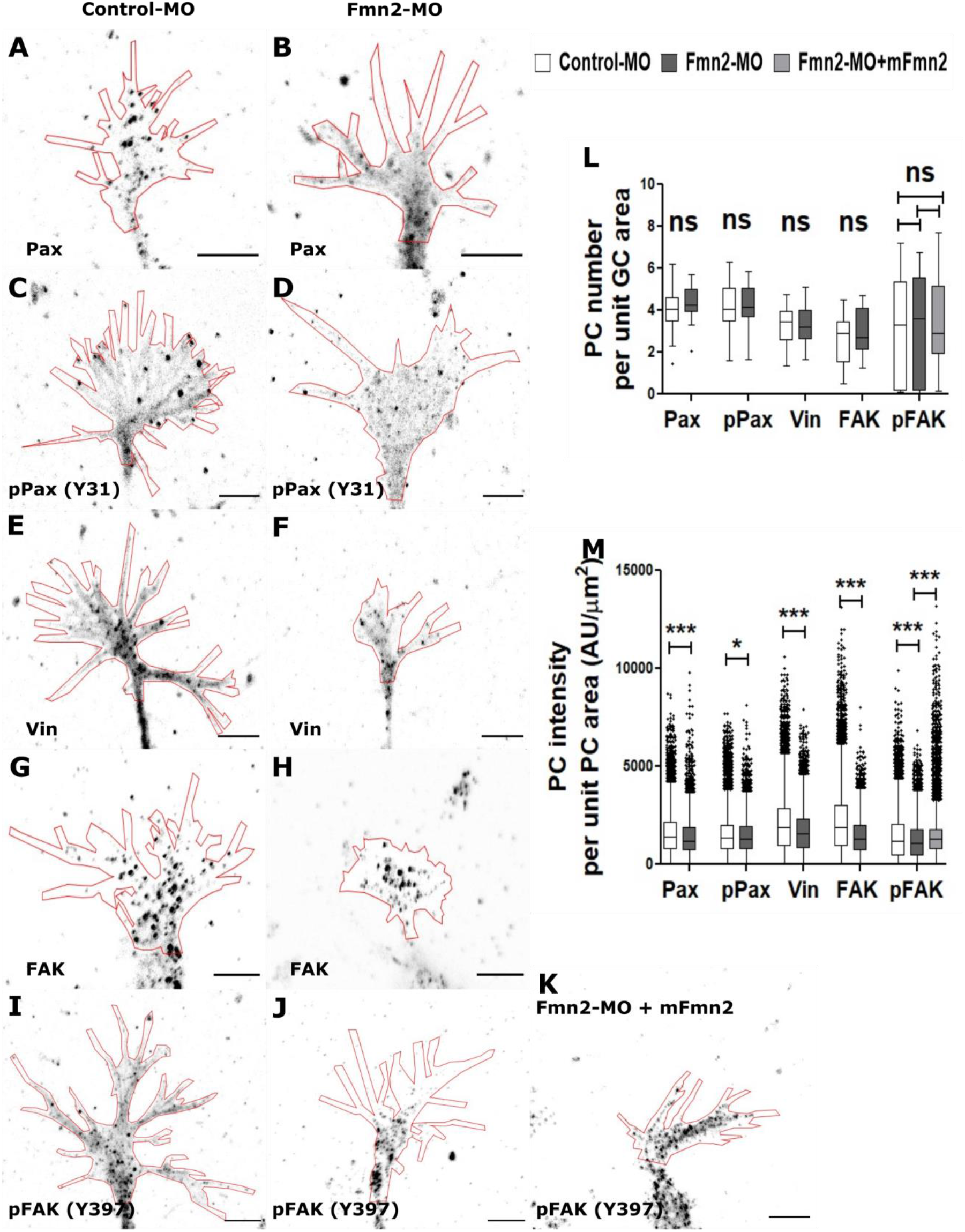
Growth cone point contact stability is regulated by Fmn2. (A and B) Representative images of spinal growth cones transfected with Control-MO or Fmn2-MO and immunostained for endogenous Paxillin (Pax). (C and D) Representative images of spinal growth cones transfected with Control-MO or Fmn2-MO followed by immunostaining for endogenous Y31 phosphorylated Paxillin (pPax (Y31)). (E and F) Representative images of spinal growth cones transfected with Control-MO or Fmn2-MO and immunostained for endogenous Vinculin (Vin). (G and H) Representative images of spinal growth cones transfected with Control-MO or Fmn2-MO and immunostained for endogenous Focal adhesion kinase (FAK). (I, J and K) Representative images of spinal growth cones transfected with Control-MO, Fmn2-MO or Fmn2-MO + mFmn2 (morpholino-resistant mouse Fmn2 cDNA) and immunostained for endogenous Y397 phosphorylated Focal adhesion kinase (pFAK (Y397)). In all micrographs (A - K), the outline of the growth cone perimeter is traced using the co-transfected GFP signal. (L) Quantification of the number of growth cone point contacts normalised to growth cone area showing no significant change upon Fmn2 depletion in all point contact markers tested. (M) Quantification of fluorescence intensity at individual point contacts (measured in arbitrary units (AU)) proteins normalised to their respective area). Number of growth cones analysed for Pax are Control-MO = 38 and Fmn2-MO = 25. Number of growth cones analysed for pPax (Y31) are Control-MO = 32 and Fmn2-MO = 19. Number of growth cones analysed for Vin are Control-MO = 33 and Fmn2-MO = 30. Number of growth cones analysed for FAK are Control-MO = 40 and Fmn2-MO = 31. Number of growth cones analysed for pFAK (Y397) are Control-MO = 53, Fmn2-MO = 48 and Fmn2-MO+mFmn2 =66. Number of point contacts detected and analysed for Pax are Control-MO = 12593 and Fmn2-MO = 9291. Number of point contacts detected and analysed for pPax (Y31) are Control-MO = 10779 and Fmn2-MO = 6498. Number of point contacts detected and analysed for Vinculin are Control-MO = 11319 and Fmn2-MO = 6394. Number of point contacts detected and analysed for FAK are Control-MO = 8278 and Fmn2-MO = 6019. Number of point contacts detected and analysed for pFAK (Y397) are Control-MO = 13540, Fmn2-MO = 9209 and Fmn2-MO+mFmn2 = 12673. Data are represented as box and whisker plots using the Tukey method. The horizontal line inside the box represents the median. Statistical comparisons were performed using the Mann-Whitney test; *, p≤0.05, **, p≤0.01, ***, p≤0.001, ns, non-significant. Scale bar, 5 μm in all images.

Upon integrin engagement, paxillin (Pax) is one of the early proteins recruited to the adhesion complex. Analysis of Pax staining of growth cones revealed that the knockdown of Fmn2 did not influence the number (normalised to the spread area of the growth cone) of Pax-labelled point contacts (Figure 2L). However, the area normalised signal intensity, reflecting the extent of recruitment of Pax to point contacts, was reduced upon reduction in Fmn2 (Figure 2M; Fmn2-MO: 1393 ± 9.83 AU/μm^2^; Control-MO: 1588 ± 9.74 AU/μm^2^).

In non-neuronal cells, phosphorylation of paxillin at tyrosine 31 (pPax) is mediated by FAK and is thought to be essential for focal adhesion assembly and lamellipodial extension (Zaidel-Bar *et al*. 2006). We investigated if pPax is affected in neurons upon loss of Fmn2. Our observations revealed that perturbation of Fmn2 leads to loss of point contacts labelled with pPax (Y31) at the central growth cone as well as filopodial tips.

Quantification of pPax further confirmed that even though the pPax-positive point contact number normalised to growth cone area remains unchanged (Figure 2L), the individual point contact intensity (Figure 2M; Fmn2-MO 1484 ± 12.00 AU/μm^2^; Control-MO: 1582 ± 11.13 AU/μm^2^) decreased after Fmn2 knockdown.

Pax interacts with vinculin (Vin) facilitating the coupling of the adhesion complex with F-actin. Like Pax, reduction of Fmn2 resulted in significant reduction of Vin-positive point contact signal intensity at individual point contacts (Figure 2M; Fmn2-MO: 1688 ± 14.09 AU/μm^2^; Control-MO: 2080 ± 13.82 AU/μm^2^) while the number of Vin-containing point contacts was unchanged (Figure 2L).

Focal adhesion kinase (FAK), a non-receptor tyrosine kinase, is activated downstream of integrin clustering and is one of the early signalling proteins recruited upon initial point contact assembly. We detected FAK in point contacts in filopodia (including filopodial tips) and throughout the growth cone area. Quantitation of the point contacts again revealed that while the number of FAK-positive point contacts was unaffected (Figure 2L) the FAK signal intensity (Figure 2M; Fmn2-MO: 1459 ± 12.41 AU/μm^2^; Control-MO: 2198 ± 18.74 AU/μm^2^) were reduced upon Fmn2 depletion.

As indicated earlier, autophosphorylation of FAK at the Y397 residue (pFAK) is an indicator of point contact stability and is central to force-dependent maturation of the point contacts (Robles and Gomez 2006; Chacon *et al*. 2012; Moore *et al*. 2012). We immunostained neurons with anti-pFAK antibody to assess the effect of Fmn2 depletion on point contact stability. We observed loss of filopodial tip adhesions labelled by pFAK (Y397) along with an overall loss of signal intensity from the central region of the growth cone. These results are consistent with the previous report from our lab (Sahasrabudhe *et al*. 2016). Fmn2 knockdown did not change the number of pFAK-containing point contacts (Figure 2L). However, reduction in individual pFAK-labelled point contact signal intensity was observed upon reduction of Fmn2 (Figure 2M; Fmn2-MO: 1223 ± 10.06 AU/μm^2^; Control-MO: 1410 ± 10.24 AU/μm^2^).

To test for sufficiency, we co-transfected morpholino-resistant mouse Fmn2 (mFmn2) along with anti-Fmn2 morpholinos and evaluated point contact function using anti-pFAK immunofluorescence (Figure 2K). Overexpression of morpholino-resistant mouse Fmn2 was able to partially rescue the reduction in the individual point contact signal intensity per unit point contact area (Figure 2M; Fmn2-MO+mFmn2 1440 ± 10.22 AU/μm^2^).

Collectively, our analysis of point contacts in growth cones revealed a common trend that suggests that the initial assembly of point contacts is unaffected (as the number of point contacts is unchanged). However, the reduction in the signal intensity of point contacts by multiple markers is suggestive of a role of Fmn2 in stabilization or maturation of these contacts. This observation is in line with our previous analysis of focal adhesion stability in fibroblasts where Fmn2 depletion enhanced focal adhesion disassembly without affecting assembly rates (Sahasrabudhe *et al*. 2016).

### Fmn2 couples F-actin retrograde flow to the substrate

The reduced growth cone translocation rates and the compromised ECM substrate attachment, as indicated by our analysis of the growth cone point contacts, in Fmn2 deficient neurons are suggestive of a compromised molecular clutch. Our previous work on fibroblasts demonstrated that Fmn2 localized to ventral actin stress fibres spanning focal adhesions. Fmn2 was juxtaposed with focal adhesion components and its localization suggested a possible role in coupling the actomyosin system to the adhesion complex (Sahasrabudhe *et al*. 2016). As loss of this coupling compromises force-dependent stabilization of adhesion structures, we evaluated the coupling of the F-actin systems directly by assessing the F-actin retrograde flow in growth cones. Reduction in F-actin retrograde flow indicates stronger engagement of the molecular clutch whereas an increase in the flow velocity would implicate a weakly engaged clutch.

Spinal neurons transfected with morpholinos and pCAG-mGFP-β-actin were subjected to live imaging to assess F-actin retrograde flow. Kymographs were generated from different regions of the growth cone, and retrograde flow velocities were computed (Figure 3A-E). These studies revealed that depletion of Fmn2 increased F-actin retrograde flow in the lamellipodial and central regions of the growth cone (Figure 3F; 0.0576 ± 0.0031 μm/sec; Control 0.0388 ± 0.0036 μm/sec).

**Figure 3:**
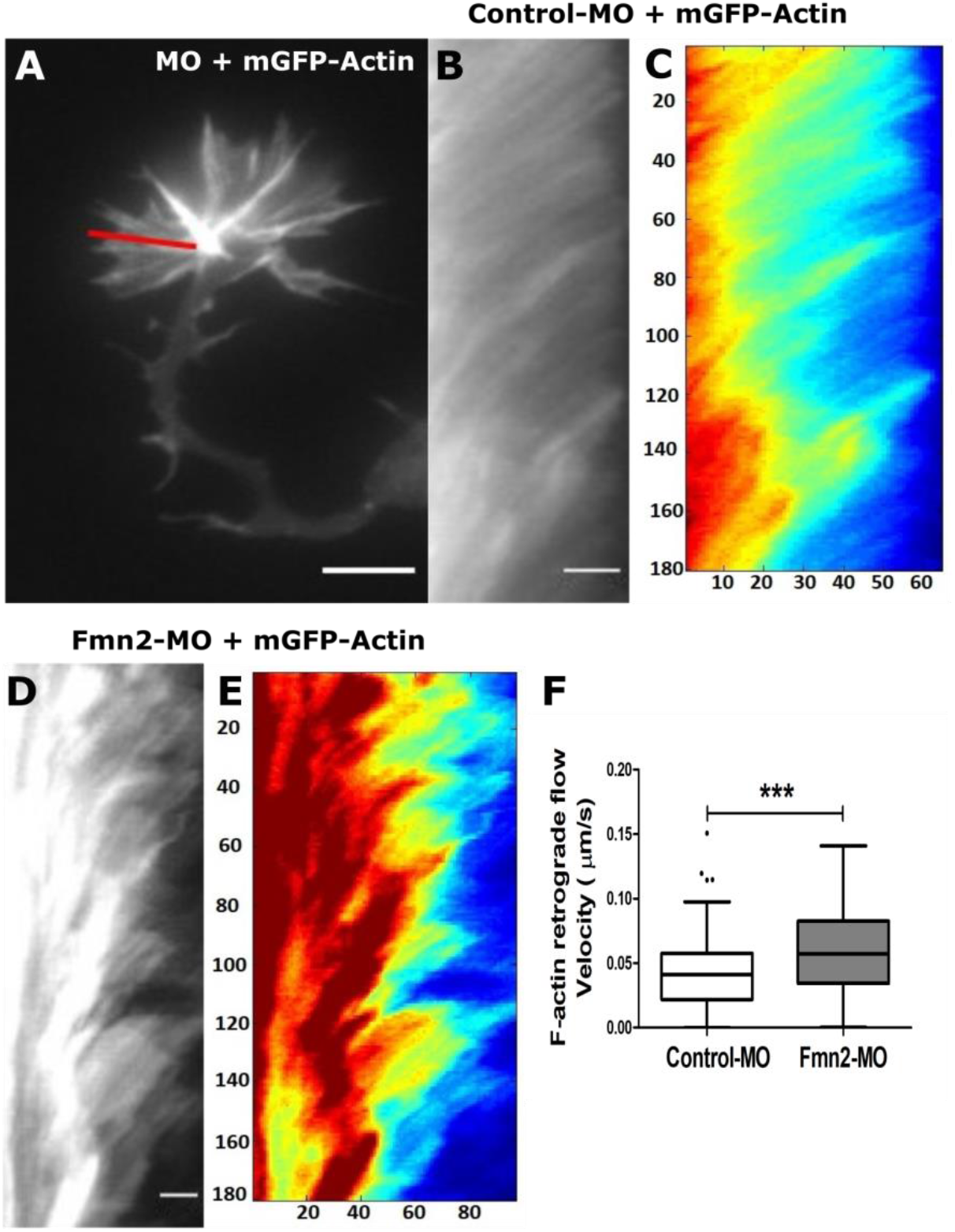
Clutch-activity of Fmn2 couples the F-actin retrograde flow to adhesion complexes. (A) Representative micrograph of a growth cone transfected with morpholinos and expressing mGFP-Actin. Kymographs were generated by drawing a line on the growth cone as shown in the figure. (B, D) Representative kymographs of growth cones transfected with Control-MO (B) or Fmn2-MO (D). (C, E) Heat maps with the x-axis representing displacement (pixels) and the y-axis representing time (seconds) were generated from kymographs (B and D) to determine the velocity of F-actin retrograde flow. Warmer colours indicated higher signal intensity. (F) Quantification of the velocity of F-actin retrograde flow. The number of growth cones analysed for Control-MO and Fmn2-MO are 28 and 23, respectively. Column graph represents the mean and the error bar represents the SEM. Statistical comparisons were performed using the Mann-Whitney test; ***, p≤0.001. Scale bar, 5 μm in (A) and 2 μm in (B, D).

The observation that loss of Fmn2 increased the rate of F-actin retrograde flow indicates a weakly engaged molecular clutch. Such a scenario could lead to compromised force-dependent stabilisation of point contacts resulting in unstable adhesion with the ECM and compromised translocation rates.

### Traction force generation by growth cones is mediated by Fmn2

Mechanical coupling between the actin cytoskeleton and the ECM at adhesion sites is necessary for the coordinated generation of traction forces onto the growth substrate and is essential for translocation. Our observation that Fmn2 deficient growth cones have destabilized adhesion complexes that are weakly coupled to the actin cytoskeleton prompted us to directly evaluate the traction forces generated by growth cones. Previous studies indicate that the traction forces generated in the central region of growth cone dominate over traction generated by filopodia (Betz *et al*. 2011; Hyland *et al*. 2014). Traction Force Microscopy (TFM) analysis was performed on morpholino and GFP transfected growth cones cultured for 48 hours on fibronectin-coated, compliant polyacrylamide gels (stiffness: 0.438 ± 0.064 kPa) embedded with fluorescent beads. Images of the growth cone and beads were acquired pre and post trypsin-induced deadhesion and analyzed using the Fourier Transform Traction Cytometry (FTTC) method (Butler *et al*. 2002) (Figure 4A, B).

**Figure 4:**
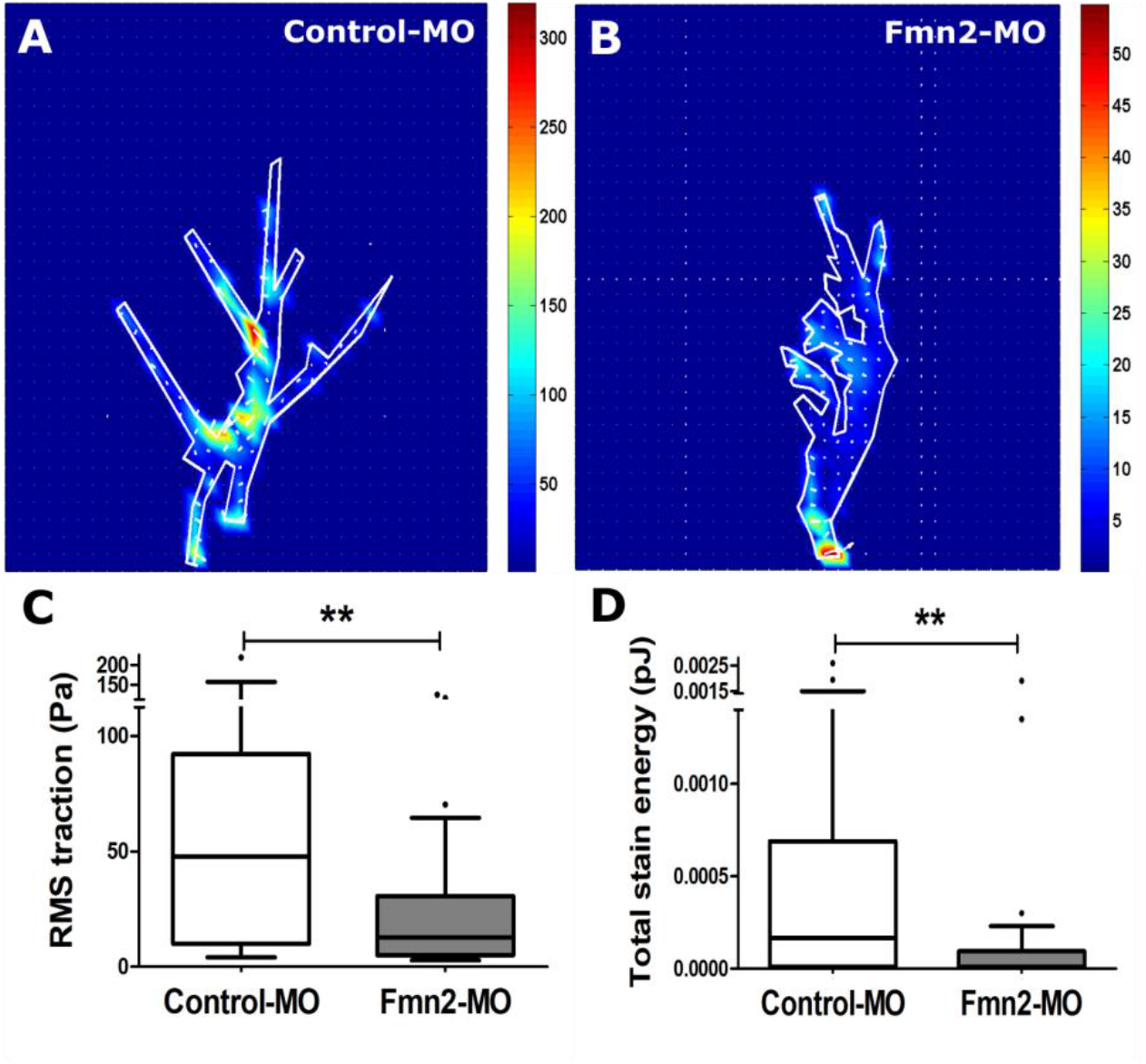
Fmn2 is required for the generation of traction forces by growth cones. (A and B) Representative heat maps of traction forces generated by Control-MO or Fmn2-MO transfected growth cones on compliant gels. Heat map scale represents traction in Pa. (C) Quantification of root mean square (RMS) traction generated growth cones reduces in growth cones lacking Fmn2 as compared to controls. (D) Total strain energy generated by growth cones is reduced in Fmn2 morphant growth cones as compared to the control growth cones. The number of growth cones analysed for Control-MO and Fmn2-MO are 36 and 38, respectively. Data are represented as box and whisker plots using the Tukey method. The horizontal line inside the box represents the median. Statistical comparisons were performed using the Mann-Whitney test; **, p≤0.01.

In spite of intrinsic variations, we found that Fmn2 depletion knockdown reduced the magnitude of the traction forces (Figure 4C; Fmn2-MO: 24.27 ± 4.885 Pa) compared to those generated by control growth cones (Figure 4C; 56.41 ± 8.530 Pa). The total strain energy was also reduced in Fmn2 depleted growth cones (Figure 4D; Fmn2-KD: 0.000132 ± 0.000060 pJ; Control 0.000448 ± 0.000102 pJ).

Collectively, our data suggest that Fmn2 mediates the coupling of the actin cytoskeleton with the adhesion complexes and, in turn, regulates point contact stability and the development of traction stresses. Compromised traction forces at point contacts result in the reduced translocation rates in Fmn2 depleted growth cones.

## DISCUSSION

In axonal growth cones, rapid assembly of actin filaments at the leading edge along with myosin activity generates a retrograde flow of F-actin. Coupling of F-actin retrograde flow with the ECM and adhesion sites allows the transmission of the actomyosin forces onto the substrate as traction stresses necessary for motility. Cell surface receptors, like the integrin receptors, bind extracellular ECM molecules while engaging with the actomyosin system via clutch molecules on the intracellular side. A number of scaffolding and signalling molecules mediate these finely-tuned interactions and form a mechanochemically regulated adhesion complex that functions as a molecular clutch. While most of the studies investigating the engagement of the actomyosin network with the ECM have been undertaken in 2-dimensional culture systems, recent work has also found evidence for clutch activities during haptotactic guidance *in vivo* (Minegishi *et al*. 2018).

We find that Fmn2 regulates growth cone translocation by functioning as a clutch molecule. Reduction of growth cone motility upon Fmn2 depletion is associated with a major remodelling of the point contacts. Our data, evaluating several different constituents of point contacts, suggest that Fmn2 does not mediate point contact initiation and assembly but is rather involved in their maintenance and stability. Knockdown of Fmn2 does not change the number of point contacts suggesting no involvement in mediating point contact assembly. However, the recruitment of the constituent point contact proteins is attenuated upon reduction of Fmn2 protein. Fmn2 may mediate either maturation of point contacts or regulate their stability. While not ruling out the former possibility, we favour the latter hypothesis for the following reasons. Autophosphorylation of FAK at tyrosine 397 is a robust readout of integrin-dependent development of stable substrate attachments and is dependent on mechanical feedback on the integrin molecules (Chacon *et al*. 2012; Robles and Gomez 2006; Moore *et al*. 2012). The depletion of Fmn2 reduces the pFAK signal and is suggestive of loss of point contact stability. We have previously shown that Fmn2 regulates tip adhesion stability and contractility of growth cone filopodia (Sahasrabudhe *et al*. 2016). Further, we have directly evaluated focal adhesion assembly and disassembly in fibroblasts and reported that knockdown of Fmn2 did not affect assembly rates but instead enhances focal adhesion disassembly (Sahasrabudhe *et al*. 2016). As reported here, the pFAK signal is also decreased in fibroblasts transfected with siRNAs against Fmn2 (Sahasrabudhe *et al*. 2016). The sensitivity to Fmn2 is observed only in cells growing on fibronectin but not when plated on glass and suggests regulatory specificity towards integrin-dependent adhesions (Figure S2). These observations in growth cones and fibroblasts strongly suggest that Fmn2 depletion affects the stability of point contacts which results in attenuated local recruitment of point contact proteins. However, it remains formally possible that Fmn2 knockdown reduces the expression point contact proteins in growth cones to limiting amounts and warrants further experimentation on neurons derived from Fmn2 knockout animals.

The molecular clutch model underscores a regulatable physical linkage of the F-actin retrograde flow with the ECM, via the adhesion complexes. Thus a strongly engaged clutch is correlated with a reduction in the centripetal F-actin flow and increased motility. We find that Fmn2 depletion results in increased F-actin retrograde flow rates. This observation suggests a weakly engaged, slippage-prone clutch activity. In non-neuronal cells, ventral stress fibres engage with focal adhesions at their two ends and the contractility of these exaggerated actin bundles is necessary for the stability of the adhesion complexes (Colombelli *et al*. 2009; Oakes *et al*. 2012; Burridge and Wittchen 2013; Chang and Kumar 2013). We suggest that the weakly engaged clutch observed upon Fmn2 reduction results in the compromised stability of the growth cone point contacts. Fmn2, due to its actin-binding abilities, localises diffusely in the central region of the growth cone and is enriched in the F-actin dense regions like the filopodia (Sahasrabudhe *et al*. 2016). While we were unable to evaluate the actin filaments engaged with the point contacts in the growth cone, in fibroblasts Fmn2 decorates ventral stress fibres and is found in close apposition to the focal adhesions (Sahasrabudhe *et al*. 2016). These observations suggest that Fmn2, directly or indirectly, couples the actomyosin system to the adhesion complex and functions as a clutch molecule.

It is also possible that Fmn2 changes the actin architecture in a manner that the generation and transmission of forces generated by actomyosin activity are compromised. However, such a scenario would likely result in a decrease in F-actin retrograde flow rates. In Fmn2 depletion, however, there is an increase in the F-actin retrograde flow with a concomitant decrease in traction stresses. In non-neuronal cells, the formin family member FHOD1 is required for assembling integrin cluster - associated actin structures (Iskratsch *et al*. 2013). Though we do not know if Fmn2, like FHOD1, induces actin assembly from integrin clusters in growth cones, there are distinct differences in their modes of action. Unlike Fmn2, depletion of FHOD1 reduces the retrograde flow of F-actin (Iskratsch *et al*. 2013). Additionally, we have previously shown that the contribution of canonical formin activities of Fmn2, like actin nucleation and F-actin filament elongation, are modest in growth cones as Fmn2 depletion does not affect either initiation or elongation of filopodia (Sahasrabudhe *et al*. 2016). Thus it appears that a major function of Fmn2 in growth cone motility is as a clutch molecule stabilizing adhesions with the ECM. This conclusion is consistent with the earlier identification of Fmn2 in myosin-dependent adhesomes in non-neuronal cells (Kuo *et al*. 2011). However, the possibility that Fmn2 indirectly destabilises point contacts via a change in actin architecture cannot be ruled out.

Traction stresses generated by the growth cone propel it forward. The physical coupling of the F-actin network with the ECM allows the transmission of the actomyosin forces as traction on the substrate. Thus, a weakly engaged clutch is expected to develop compromised traction forces. Direct evaluation of traction forces using TFM revealed significantly attenuated traction stresses. The strain energy, the total energy transferred from the cell to the substrate, was also reduced and underscored the compromised force transmission from the growth cone onto the ECM.

Taken together, our observations suggest that Fmn2 regulates the clutch activity at adhesion sites and thereby mediates growth cone translocation. For haptotactic responses of growth cones, the local differences in clutch activity in response to external cues are believed to steer growth cones and impart directionality (Baba *et al*. 2018). In *Xenopus* spinal neurons, local changes in F-actin retrograde flow at point contacts have been shown to mediate axon guidance (Santiago-Medina *et al*. 2013). As regulatory modalities of Fmn2 are unknown, it remains to be investigated if cue-dependent regulation of Fmn2 clutch activity can drive growth cone turning.

Fmn2 is implicated in several neurodevelopmental and neurodegenerative disorders in both humans and rodent models (Peleg *et al*. 2010; Almuqbil *et al*. 2013; Law *et al*. 2014; Agís-Balboa *et al*. 2017; Marco *et al*. 2018). However, the molecular basis of Fmn2 function in neurons is largely unknown. The Fmn2-dependent mechanisms identified in this study are likely to contribute to the development of a mechanistic framework for Fmn2 function in neurons. Further, Fmn2 is known to be de-regulated in a number of cancers (Oncomine and The Cancer Genome Atlas (TCGA) public databases). Findings reported here may be of significance to cancer progression, especially in the context of cancer cell - ECM associations and metastasis.

In summary, our work identifies Fmn2 as a novel clutch molecule mediating growth cone motility. This function is likely to be a major contributor to the developmental role of Fmn2 in regulating axon outgrowth and guidance. Our study also identifies a novel function for formins and highlights the functional diversity of this family of proteins in cytoskeletal remodelling.

## ETHICS APPROVAL

All protocols used in this study were approved by the Institutional Animal Ethics Committee and the Institutional Biosafety Committee of IISER Pune.

## AVAILABILITY OF DATA AND MATERIAL

All data generated or analysed during this study are included in this published article. The raw data are available from the corresponding author on a reasonable request.

## AUTHOR CONTRIBUTIONS

Conceptualization, K.G., S.P.M. and A.G.; Investigation and analysis, K.G. (all experiments), S.P.M. (optimizing traction force microscopy and methods to analyse F-actin retrograde flow; preparation of polyacrylamide gels for AFM analysis), K.L.S. / S.S. (AFM analysis of polyacrylamide gels); Writing, K.G. and A.G.; Funding Acquisition, A.G.. All authors gave final approval for publication and agree to be held accountable for the work performed therein.

## ACKNOWLEDGEMENTS

The authors acknowledge the IISER Pune Microscopy Facility at IISER Pune for access to equipment and infrastructure. The authors thank Prof. N. K. Subhedar, IISER Pune for his critical reading and suggestions on early drafts of this manuscript.

