## Supplementary Information for "Fmn2 regulates growth cone motility by mediating a molecular clutch to generate traction forces"

**This file includes:**

Figures S1 to S2
Legend for Movie S1
Supplementary Materials and Methods
Supplementary References

**Other supplementary materials for this manuscript include the following:**

Movie S1

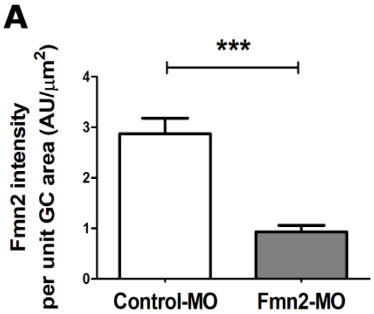


**Figure S1: Fmn2 morpholino robustly reduces endogenous Fmn2 protein levels**

(A) Quantification of fluorescence intensity from endogenous Fmn2 in chick spinal neuron growth cones (measured in arbitrary units (AU)) transfected with Control-MO (n = 24) or Fmn2-MO (n = 16). Data are represented as box and whisker plots using the Tukey method. The horizontal line inside the box represents the median. Statistical comparisons were done using Mann-Whitney test; ***, p≤0.001).

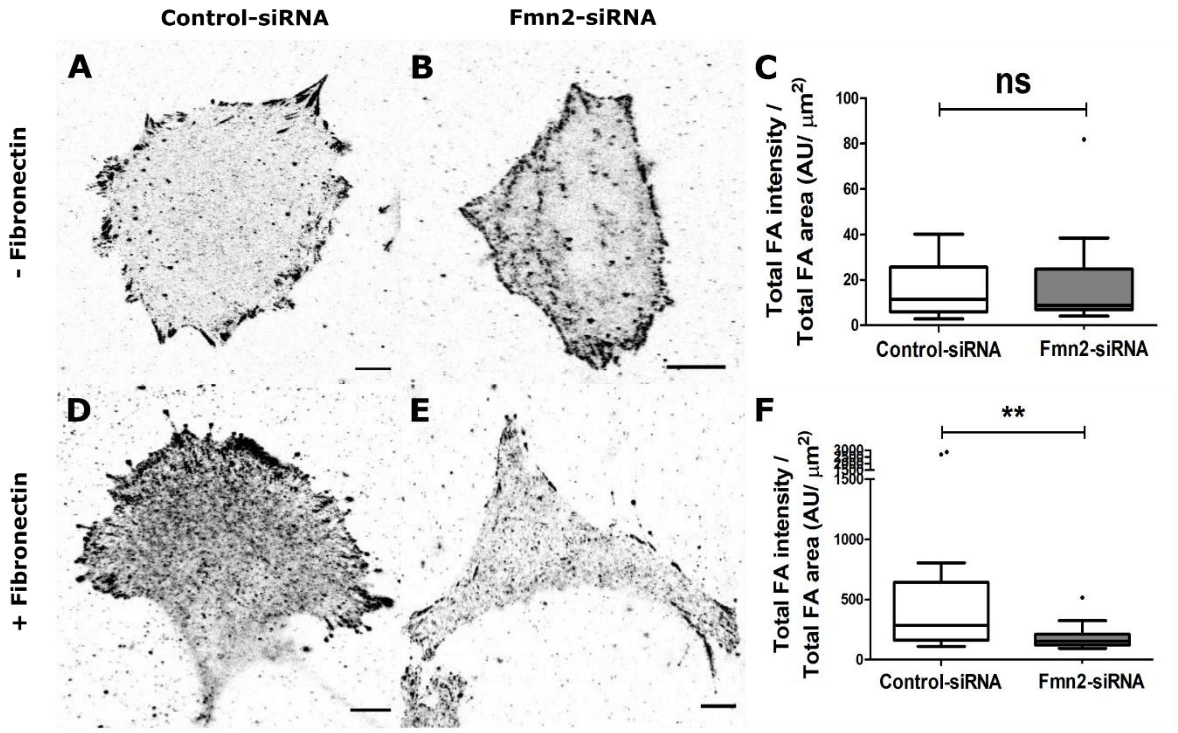


**Figure S2: Fmn2 regulates focal adhesion stability in fibroblasts only when plated on fibronectin.**

(A and B) Representative micrographs of Control-siRNA or Fmn2-siRNA transfected NIH3T3 fibroblasts cultured on fibronectin-coated substrate and subsequently immunolabelled with pFAK (Y397). (C) Quantification of pFAK (Y397) immunofluorescence at focal adhesions (measured in arbitrary units (AU)) normalised to total FA area. (D and E) Representative micrographs of Control-siRNA or Fmn2-siRNA transfected NIH3T3 fibroblasts cultured on fibronectin-coated substrate and subsequently immunolabelled with pFAK (Y397). (F) Quantification of pFAK (Y397) immunofluorescence at focal adhesions (measured in arbitrary units (AU)) normalised to total FA area. Number of cells cultured without fibronectin and analysed for both Control-siRNA and Fmn2-siRNA are both 22. Number of cells cultured on fibronectin and analysed for control-siRNA and Fmn2-siRNA are 19 and 20, respectively. Data are represented as box and whisker plots using the Tukey method. The horizontal line inside the box represents the median. Statistical comparisons were done using Mann-Whitney test; **, $p \leq 0.01$.

**LEGEND FOR SUPPLEMENTARY MOVIE**

**Movie S1: Fmn2 regulates growth cone translocation**

**Representative timelapse video of translocation of spinal neuron growth cones** transfected with Control-MO (top), Fmn2-MO (middle) or Fmn2-MO + mFmn2 (bottom). mFmn2, morpholino-resistant mouse Fmn2. The morpholinos were co-transfected with GFP to identify transfected neurons. Scale bar, 5 µm; time stamp, min:sec.

### SUPPLEMENTARY MATERIALS AND METHODS

#### Fmn2 immunofluorescence in growth cones

Spinal neurons were electroporated and cultured with Control and Fmn2 morpholinos (as indicated in the Material and Methods section) for 24 hrs and fixed using 4% PFA (Electron microscopy sciences) and 0.05% glutaraldehyde (Electron microscopy sciences) in PBS (mentioned earlier) for 10 min at room temperature. After removing PBS, cells were washed with PBS three times for 10 mins each. The cells were permeabilized using PBST (0.1% Triton-X in PBS) for 30 min. After permeabilization, two brief PBS washes were given and the cells were blocked in 3% BSA for 60 min at room temperature. The blocking solution was replaced by antibody against chick Fmn2 (developed in house and characterized in (Sahasrabudhe *et al.* 2016) in 1:200 dilution in 3% BSA and incubated at $4^0$C for 14-16 hrs. Subsequently, after removing primary antibody, the cells were washed four times with PBS + 0.1% TritonX, for 10 minutes each. Later the sample was incubated at room temperature for 60 mins with secondary antibody (Invitrogen) diluted 1:1000 in 3% PBS. After removing secondary antibody, the cells were washed again in PBST four times, 10 min each. The cells were further labelled with phalloidin (Invitrogen) diluted 1:200 in blocking for 30-45 mins at room temperature. Finally, the cells were washed thrice in PBST for 10 min each and mounted in 80% glycerol.

#### NIH3T3 culture, transfection and pFAK (Y 397) staining

NIH3T3 mouse embryo fibroblast cell line was procured from Dr. N. Balasubramanian (IISER, Pune) and checked for contamination prior to use. Cells at passage numbers 25 to 40 were used for experiments. Cells frozen previously in liquid nitrogen were thawed and cultured in complete DMEM (Lonza) with 1x PenStrep (Gibco), 1x sodium pyruvate (Invitrogen) and 10% heat inactivated FBS at $37^0$C with 5% $CO_2$. After the cells were 80-90% confluent, the media was removed and cells were washed with DPBS without $Ca^{2+}$ and $Mg^{2+}$ (Lonza), thereafter 1-2 ml of 1x trypsin-EDTA was added (Lonza). Following trypsinization, approximately 3 x $10^5$ cells were re-seeded in 10 ml of complete DMEM.

2 x $10^4$ NIH3T3 cells were seeded on glass bottom chambers (Labtek) with or without pre-coating the chambers with 20µg/ml fibronectin (Sigma) in PBS for 1 hour at $37^0$C. The cells were allowed to adhere for 14-16 hrs and then transfected using Lipofectamine 2000 (Invitrogen). After cells were about 60-70% confluent, the cell culture media was replaced by OptiMEM (Gibco) and cells were incubated at $37^0$C for 20 min. Simultaneously the transfection mix was prepared with OptiMEM, 1-5µg Paxillin-GFP plasmid DNA, 3-5µl lipofectamine and control LacZ siRNA (Control-siRNA; sense strand: CGUCGACGGAAUACUUCGAUU) or mouse Fmn2 siRNA (Fmn2-siRNA; sense strand: UGGUUAGACUUGUGGGUAAUU) and incubated at room temperature for 20 mins. Subsequently, the transfection mix was added onto the cells and they are incubated for 4 hours at $37^0$C after which the media was supplemented by complete DMEM and 30% FBS.

24 hrs post-transfection, NIH3T3 cells were fixed using 4% PFA (Electron microscopy sciences) and 0.05% glutaraldehyde (Electron microscopy sciences) in PBS for 10 min at room temperature. After removing PBS, cells were washed with PBS three times for 10 mins each. The cells were permeabilized using 0.1% Triton-X in PBS for 30 mins. After permeabilization, two brief PBS washes were given and the cells were blocked in 3% BSA for 60 min at room temperature. The blocking solution was replaced by antibody against phosphor-FAK (Y 397) (Abcam) in 1:1000 dilution in 3% BSA and incubated at $4^0$C for $1^4$-16 hrs. Subsequently after removing primary antibody, the cells were washed four times with PBS + 0.1% TritonX, 10 times each. Later the sample was incubated at room temperature for 60 min with secondary antibody (Invitrogen) diluted 1:1000 in 3% PBS. After removing secondary antibody, the cells were washed again in PBST four times, 10 min each. Subsequently, the cells were labelled with phalloidin (Invitrogen) diluted 1:200 in blocking for 30-45 mins at room temperature. Lastly the cells were washed thrice in PBST, 10 min each and mounted in 80% glycerol.

Spinal neurons labelled with anti-Fmn2 antibody were imaged using PlanApo 60x/1.4 oil immersion objective on a Leica Sp8 confocal system. All the imaging conditions were kept identical for imaging Control and Fmn2 morpholino electroporated neuronal cultures to analyse intensity changes. For imaging immunofluorescence of pFAK, PlanApo 63x/1.40 oil immersion objective was used on Zeiss 710 inverted confocal microscope.

#### Analysis and data representation

To analyse endogenous Fmn2 levels in the growth cones, the outline of growth cones was manually drawn in Fiji based on cytoplasmic GFP channel. This ROI was overlaid on anti-Fmn2 channel and the mean grey values were determined. Mean grey values normalised to respective growth cone areas were plotted in GraphPad Prizm 5.0 and compared using Mann-Whitney test in Prism 5.

To assess NIH3T3 cells for pFAK intensity, the paxillin expression images were uploaded on Focal adhesion analysis server (FAAS) (Berginski and Gomez 2013) to get masks based on focal adhesions marked by paxillin. This mask was overlaid on the pFAK channel to extract the fluorescence intensities only at the demarcated focal adhesions. The intensity per unit area was plotted as Box and Whiskers (Tukey) plots and analysed using GraphPad Prism 5.
